# Low prevalence of HLA-G antibodies in lung transplant patients detected by MAIPA adapted protocol

**DOI:** 10.1101/2023.08.04.551968

**Authors:** Pascal Pedini, Lucas Hubert, Federico Carlini, Jean Baptiste Baudey, Audrey Tous, Francois Jordier, Agnès Basire, Claude Bagnis, Martine Reynaud-Gaubert, Benjamin Coiffard, Jacques Chiaroni, Monique Silvy, Christophe Picard

## Abstract

Lung transplantation is often complicated by acute and/or chronic rejection leading to graft function loss. In addition to the HLA donor-specific antibodies (HLA-DSA), a few autoantibodies are correlated with the occurrence of these complications. Recently, antibodies directed against non-classical HLA molecules, HLA-G, -E, and -F have been detected in autoimmune diseases, like systemic lupus erythematosus. Non-classical HLA molecules are crucial in the immunological acceptance of the lung graft, and some of their isoforms, like HLA-G*01:04 and -G*01:06, are associated with a negative clinical outcome. The aim of this study is to determine the frequency of detection of HLA-G antibodies in lung transplant recipients (LTRs) and their impact on the occurrence of clinical complications. After incubating the cell lines SPI-801, with and without 3 different HLA-G isoforms expression, with sera from 90 healthy blood donors and 35 LTRs (before and after transplantation), HLA-G reactivity was revealed by using reagents from commercial monoclonal antibody immobilization of platelet antigen assay (MAIPA ApDIA®). Only one serum from one blood donor had specific reactivity against the HLA-G transduced lines. Non-specific reactivity in many sera from LTRs was observed with transduced and wild type cell lines, which may suggest recognition of an autoantigen expressed by the SPI-801 cell line. In conclusion, this study allowed the development of a specific detection tool for non-denatured HLA-G antibodies. These antibodies seem uncommon, both in healthy subjects and in complicated LTRs. This study should be extended to patients suffering from autoimmune diseases as well as kidney and heart transplant recipients.

## INTRODUCTION

Lung transplantation (LT) is a therapeutic option for chronic, irreversible respiratory failure. Survival after LT has steadily increased over time, but the incidence of mild and long-term morbidity and lung dysfunction remains very high due to acute or chronic rejection (1). These clinical complications may be linked to immune dysregulation implying humoral and cellular autoimmunity and/or alloimmunity. Recent works have suggested that autoimmune reactivity is a key element in the occurrence of chronic graft rejection (2–4). Antibodies directed against Tubulin K-1, a membrane protein expressed by bronchial epithelial cells, and against collagen type V, an extracellular matrix protein normally sequestered in bronchial tissue, are strongly correlated with the occurrence of bronchiolitis obliterans syndrome (BOS) after LT (5). This production of autoantibodies may be secondary to the exposure of cryptic antigens of the self following tissue remodeling after LT, to the expression of neo-antigens in an inflammatory context, or to the stimulation of cross-reaction by antigenic mimicry between pathogens and self-antigens (6). More recently, it has been shown that the release of graft exosomes in an inflammatory context increases the risk of autoimmunity (7).

The HLA-G molecule has been suggested as a prognostic biomarker of LT outcome by two independent studies (8,9). HLA-G has both humoral and cellular anti-inflammatory immunosuppressive properties, in particular by inhibiting NK and cytotoxic T lymphocyte (CTL)-mediated activity as well as B cell activation via their inhibitory receptor (ILT-2, -4, and KIR2DL4) (10). This non-classical HLA class I molecule also acts indirectly on immune control as HLA-E preferentially loads its signal peptide (11,12). HLA-G anti-inflammatory potential is supported by its increased expression through anti-inflammatory cytokines, as IL-10, in a positive feedback mechanism, by its increased expression in autoimmune diseases, and by its involvement in promoting viral or parasitic infections escape (10).

Interestingly, HLA-G is expressed at bronchial cells membranes after interferon-β stimulation (13). Moreover, membrane-bound HLA-G expression in lung biopsies and sHLA-G expression in bronchial alveolar fluid, but not in serum, were higher in clinically stable lung transplant recipients (LTRs) than in those developing acute rejection (8,13,14).

One hundred and seventeen HLA-G alleles are currently identified, and five protein isoforms (HLA-G*01:01, *01:03, *01:04, *01:05N, 01:06) are found with a frequency > 5% in all populations (15,16). Our team and others have shown that these alleles are associated with differential sHLA-G levels (15,17,18).

HLA-G alleles have also been associated with clinical outcomes: the HLA-G*01:06∼UTR2 haplotype has been correlated with the pejorative evolution of cystic fibrosis, and the HLA-G*01:04∼UTR3 haplotype with an increase of chronic rejection, production of HLA antibodies, and a decrease in patient survival (9).

Despite its low diversity and because of its epithelial expression, HLA-G may elicit immunization mechanisms like those observed for other HLA class I molecules.

Anti-HLA-G antibodies were recently detected in patients with systemic lupus erythematosus (SLE) in a study conducted on 69 German and 29 Mexican SLE patients and 17 German healthy individuals (19,20). Multiplex Luminex®-based flow cytometry was used to screen sera for antibodies directed against non-classical HLA (HLA-G, -E, and –F) and β2m. Interestingly, anti-HLA-G IgG antibodies were detected in 30% of healthy subjects (6/17) and were more frequent than those directed against HLA-E and HLA-F. In addition, anti-HLA-G antibody levels were stable in 5 subjects over 6 months.

Our hypothesis is that HLA-G antibodies produced following LT may interfere with the HLA-G receptors and disinhibit the cellular response directed against the lung graft. Such a mechanism would participate in a persistent inflammatory response and eventually lead to respiratory failure.

The main goals of this study are 1) to detect HLA-G antibodies expressed in serum using an adapted MAIPA ApDia® kit and 2) to evaluate their impact on the occurrence of clinical complications in 35 LTRs.

## MATERIALS AND METHODS

### Samples

Two hundred and thirty-four samples were analyzed from healthy donors (N=90) and LTx patients (N=35). Sera from blood donors were collected after the medical interview and before blood donation. Sera from LTRs were collected serially before transplantation and then at 15 days (D15), 1 month (M1), 3 months (M3), and 12 months (M12) after transplantation. All LTRs were genotyped for classical HLA (HLA-A, -B, -C, -DRB1, -DQB1, and –DPB1) by NGS Omixon protocol (Omixon Biocomputing Ltd, Hungary) and for HLA-G by NG-MIX (EFS, Saint-Denis, France).

Blood donations were collected in the “Etablissement Francais du Sang”, in accordance with BSL-2 practices. A medical interview was carried out prior to blood donation to exclude donors with medical contraindications. This study was carried out in accordance with the French Public Health Code (art L1221-1), approved by institutional ethics committee and conducted in compliance with the Good Clinical Practice Guidelines, declaration of Helsinki and Istanbul. All Lung Tranplant Recipeints from the French cohort (COLT, Cohort in Lung Transplantation, l’Institut du Thorax, INSERMUMR1087/CNRS UMR 6291, CNIL 911142) were recruited in this study and gave their written informed consent to participate to the study in accordance with the Declaration of Helsinki.

These are patients sampled between 2009 and 2014, with clinical data collected in 2017.

### HLA-G and HLA-G K562 cell line transduction and expression assessment

In order to cover 81% of HLA-G haplotypes (16), cDNA coding for HLA-G*01:01 (IMGT/HLA<u>00939),</u> -G*01:04 (IMGT/HLA<u>00949</u>), -G*01:06 (IMGT/HLA<u>01357</u>) were obtained from Life Technologies (France) and cloned into the pWPXL lentiviral vector. The vector pWPXL-EGFP was carried out as a control. Lentiviral particles were generated using HEK 293T cells at 80% confluence (DSMZ, Brunswick) co-transfected with each lentiviral vector pWPXL, the vesicular stomatitis virus-G-encoding plasmid pMDG and the packaging plasmid pCMVΔR8.91 (21). Lentiviral particles were collected from day 2 (D2) to 4 (D4) and concentrated in 10% polyethylene glycol. SPI-801 cells (5.104) derived from K562 cells (DSMZ ACC-86, Brunswick, Germany) were transduced using each lentiviral particle. The same protocol was used to obtain cells expressing HLA-E*01:01 (IMGT/HLA00934) and HLA-E*01 :03 (IMGT/HLA00936).

Non-infected (WT) and transduced SPI-801 cells were screened for HLA-G and HLA-E expression by western blot and flow cytometry (Supplementary Figure 1). Western blot was performed with 4H84 antibody (HLA-G, R&D system #MABF2169). Flow cytometry was performed using a Myltenyi VYB cytometer with MEM-G/9 antibody (anti-HLA-G antibody ; Thermo Fisher #MA1-19014), 3D12 (anti-HLA-E antibody ; Thermo Fisher # 4-9953-82) or the isotype-matched control Ab (Supplementary Figure 2).

### HLA-G antibodies detection

HLA-G antibodies were screened using an adapted MAIPA protocol in accordance with the MAIPA ApDIA® kit. This kit allows the detection and identification of anti-platelet glycoprotein autoantibodies or anti-HPA (human platelet antigen) alloantibodies in serum using a platelet panel and platelet glycoprotein antigen immobilized by a monoclonal antibody. For the purpose of the present study, all MAIPA ApDIA® kit reagents were used, except for the platelet panel and platelet glycoprotein antigen, which were replaced by HLA-G transduced cell lines and WT.

Briefly, the adapted assay (Figure 1A) consisted of two consecutive incubations: sera with HLA-G transduced cell line and then with MEM-G/9 monoclonal antibody. Cells were lysed and solubilized complexes were immobilized on a plate coated with anti-mouse antibody. Complexes were revealed by immunoassay colorimetry.

**Figure 1:**
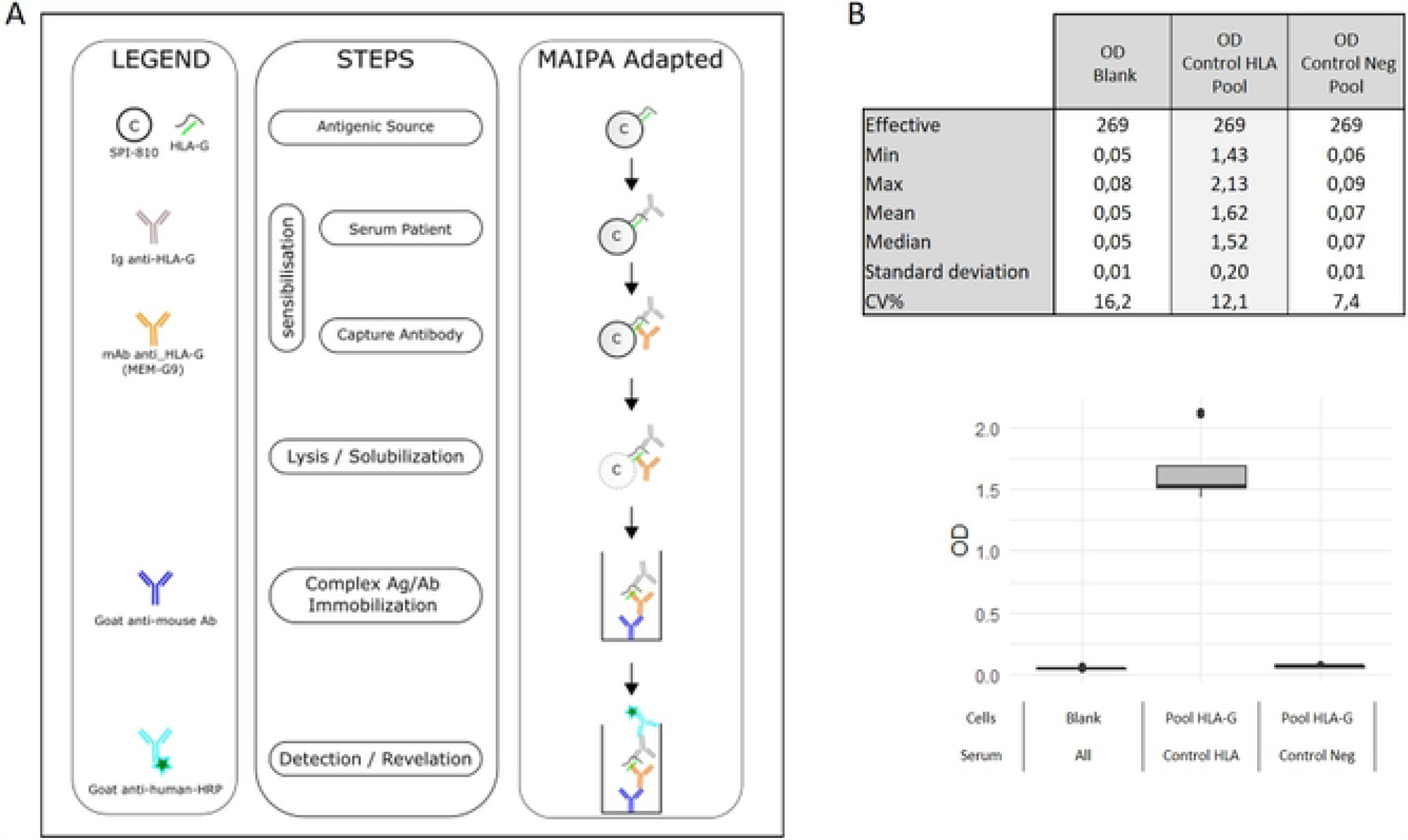
(A) Illustration of the different technical steps, adaptations and compounds for MAIPA Adapted. (B) Characteristics of MAIPA Adapted quality controls.

Serum HLA-G antibodies were detected with a pool of the three HLA-G transduced and WT cells and identified with each HLA-G transduced cell and WT cell. Negative control (Contr Neg), positive control (Contr HLA), both supplied with the MAIPA ApDIA® kit, and two blank controls were included in each assay (Figure 1B).

Detection and identification protocol consisted of incubation (30 ± 5 minutes at 36 ± 1 °C) of transduced or WT cells (N=1 × 10^6^) washed and adjusted in each well in PBS / 1% BSA / 0.33% EDTA with positive control (Contr HLA) or negative control (Contr Neg), (50μL dilution 1/1000) or with serum (50μL dilution 1/5). Microplates were centrifuged (1000g, 3 minutes) and each well was washed three times (ELISA Wash Buffer). MEM-G/9 monoclonal antibody was added (50 μg) and incubated for 30 ± 5 minutes at 36 ± 1 °C.

Cells were washed as described above and lysed (Platelet Lysis Buffer containing Triton, 130μL, 15 minutes at 2 - 8 °C). Cell lysis supernatant (100 μL) containing the solubilized captured-complex (MEM-G9 / HLA-G / sera or control anti-HLA-G) were centrifuged and were incubated in Goat anti-Mouse IgG coated microplate (30 ± 5 minutes at 36 ± 1 °C). Plates were washed as described above, and immobilized complexes were revealed by peroxidase reaction (substrate solution Chrom, 15 minutes at 36 ± 1°C in the dark). The reaction was stopped (Stop Solution, 100 μL) and read by optical density (OD) using a spectrophotometer (450 nm with reference filter 650 nm).

The assay was validated when OD values were below 0.1 for the negative control (Contr Neg) and above 2 for the positive control (Contr HLA). The cut-off OD value was set at 0.15 (Contr Neg mean OD values adjusted with blank plus 3xStandard Deviation). OD values above the cut-off were considered positive. The signal-to-noise (S/N) ratio was calculated by dividing the OD value of HLA-G or HLA-E wells by that of WT cells. Serum was considered positive for HLA-G antibodies presence when the S/N ratio was >1.5.

Screening assays were performed for each serum. Identification was performed for sera with HLA-G screening pool OD values > 0.150 and S/N ratio > 1.5.

## RESULTS

### Population characteristics

Ninety sera were collected from healthy blood donors (median age = 40 years [18–65]; sex ratio = 0.54) with no history of transfusions, organ transplantation, or recent infections. Thirty out of 41 women (73%) had at least one pregnancy.

All lung transplant recipients (median age: 40 years [19-66]; sex ratio = 0.51) underwent LTx at the Marseille Lung Transplant Center. They received a first LT (8 single LTx; 26 bilateral LTx) for cystic fibrosis (49%), emphysema (20%), pulmonary fibrosis (26%), or another diagnosis (5%). One hundred and forty-four sera were collected from 35 lung transplant recipients before and after transplant (at day 0 (D0) N=30, at day 15 (D15) N=25, at one month (M1) N=28, at three months (M3) N=32, and at twelve months (M12) N=29). Seven LTRs had acute rejection, and 7 developed chronic rejection in the first year. Seventeen produced Donor Specific Antibody (DSA) after LTx. One recipient had an acute rejection associated with DSA detection. Population characteristics are summarized in Table 1.

**Table 1:** Clinical and biological characteristics of all cohort LTRs, LTRs with at least one positive serum detected on transduced and non-transduced cell lines, and Healthy subjects (LTR : lung transplant recipient, DSA : donor specific antibody).

|  | Cohort<br>LTRs | LTRs<br>with positive serum | Healthy<br>blood |
| --- | --- | --- | --- |
| Recipient/donor |  |  |  |
| Number | 35 | 12 | 90 |
| Age |  |  |  |
| mean | 40 | 36,5 | 39,5 |
| min-max | 19-66 | 21-66 | 18-69 |
| < 40 years | 18 | 7 | 53 |
| ≥ 40 years | 17 | 5 | 37 |
| Gender |  |  |  |
| M | 17 | 5 | 41 |
| F | 18 | 7 | 49 |
| Pregnancy | ND | ND | 30 |
| First pregnancy | ND | ND |  |
| >4 pregnancies | ND | ND |  |
| Blood Group |  |  |  |
| A | 12 | 4 | 35 |
| B | 7 | 3 | 7 |
| AB | 2 | 2 | 8 |
| O | 14 | 3 | 40 |
| Pathology |  |  |  |
| Emphysema | 7 | 3 |  |
| Cystic Fibrosis | 17 | 5 |  |
| Fibrosis | 9 | 3 |  |
| Bronchial dysplasia | 1 | 1 |  |
| Histocytosis | 1 | 0 |  |
| Type of LTx |  |  |  |
| Single LTx | 9 | 3 |  |
| Bilateral LTx | 26 | 9 |  |
| Mismatch HLA |  |  |  |
| 0 | 5 | 2 |  |
| 1 – 4 | 2 | 1 |  |
| 5 – 8 | 28 | 9 |  |
| CMV Status |  |  |  |
| Absence mismatch | 25 | 8 |  |
| D+ et R- without seroconversion | 1 | 0 |  |
| D+ et R- with seroconversion | 4 | 1 |  |
| D- et R+ | 5 | 3 |  |
| Type of rejection |  |  |  |
| Acute (DSA) | 7 (7) | 2 (2) |  |
| Chronic (DSA) | 7 (6) | 1 (1) |  |
| HLA antibodies |  |  |  |
| DSA after LTx | 26 | 8 |  |
| Ab No DSA after LTx | 11 | 5 |  |
| D0 | 2 | 1 |  |
| M1 | 5 | 2 |  |
| M3 | 6 | 2 |  |
| M12 | 2 | 0 |  |
| HLA-G genotyping |  |  |  |
| G*01:01,*-- | 21 | 6 |  |
| G*01:01,G*01:06 | 4 | 3 |  |
| G*01:01,G*01:04 | 6 | 2 |  |
| G*01:01,G*01:05N | 2 | 0 |  |
| G*01:01,G*01:03 | 1 | 0 |  |
| G*01:03,G*01:06 | 1 | 1 |  |

### HLA-G cell lines transduction

The specific expression of the HLA-G isoform for each transduced cell line was assessed by western blot using 4H84 antibody and by flow cytometry using MEM/G9 antibody (Figure 2). Flow cytometry showed specific HLA-G membrane expression in HLA-G transduced cells, and no signal was detected with the MEM/G9 in transduced WT cells.

**Figure 2:**
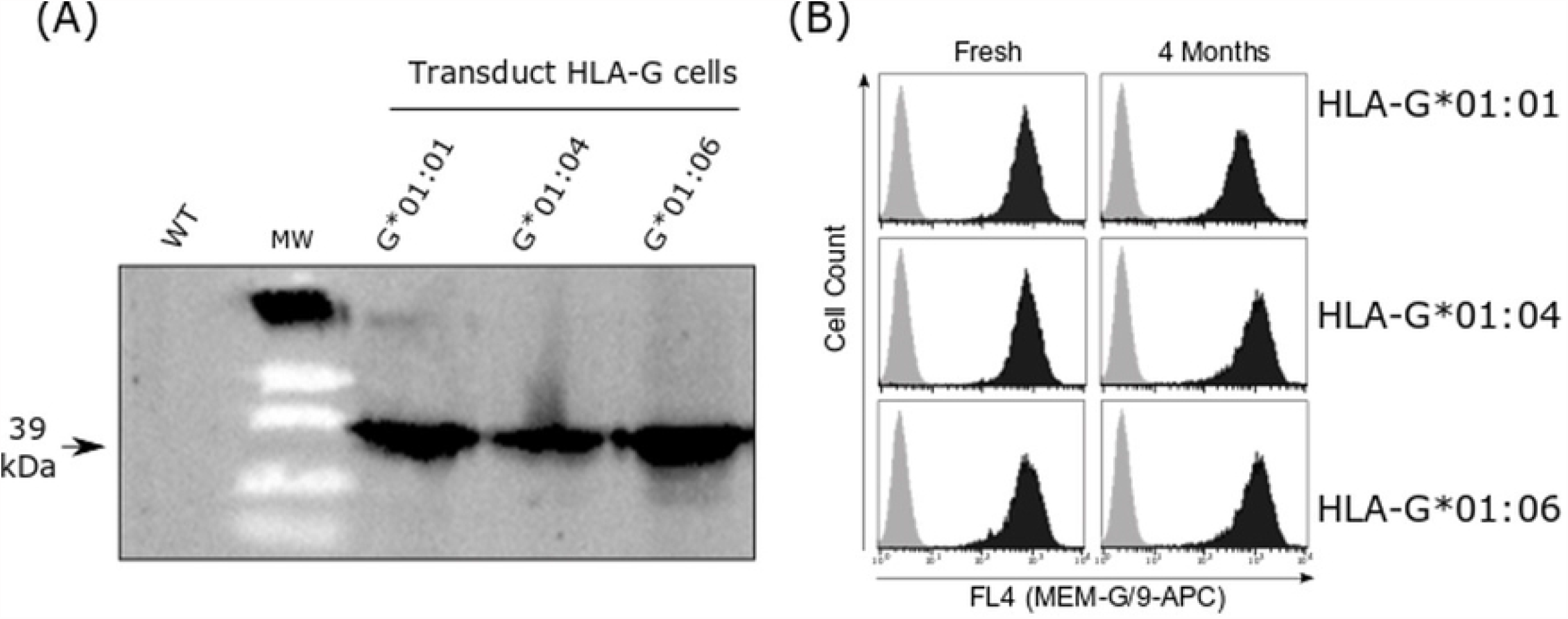
Expression of HLA-G isoform on SPI801 cell lines. (A) Western Blot analysis of HLA-G in the transduced cells in comparison with non-transduced cells line (WT). (B) Flow cytometry results showing stable expression levels of HLA-G isoforms in transduced cells lines (black) and non-transduced cells line (gray).

### HLA-G antibody detection in healthy donors

Sera from 90 healthy donors were tested (Table 2). The screening pool ODs mean was 0.07 +/-0.012 for donors, with no statistical difference between men and women (0.071 +/-0.021 vs. 0.074 +/-0.023; p = 0.45). The reactivity of the sera against the WT cell line was on average 0.066 +/-0.02, corresponding to an OD ratio mean between the HLA-G pool screening and the WT cell line of 1.02. One serum was positive for HLA-G antibody detection: a 22-year-old woman with A+ blood group. A 30-year-old man, with A+ blood group, displayed a pool reactivity OD > 150 with a ratio close to 1 between the pool and the WT cell line (Table 2).

**Table 2:**
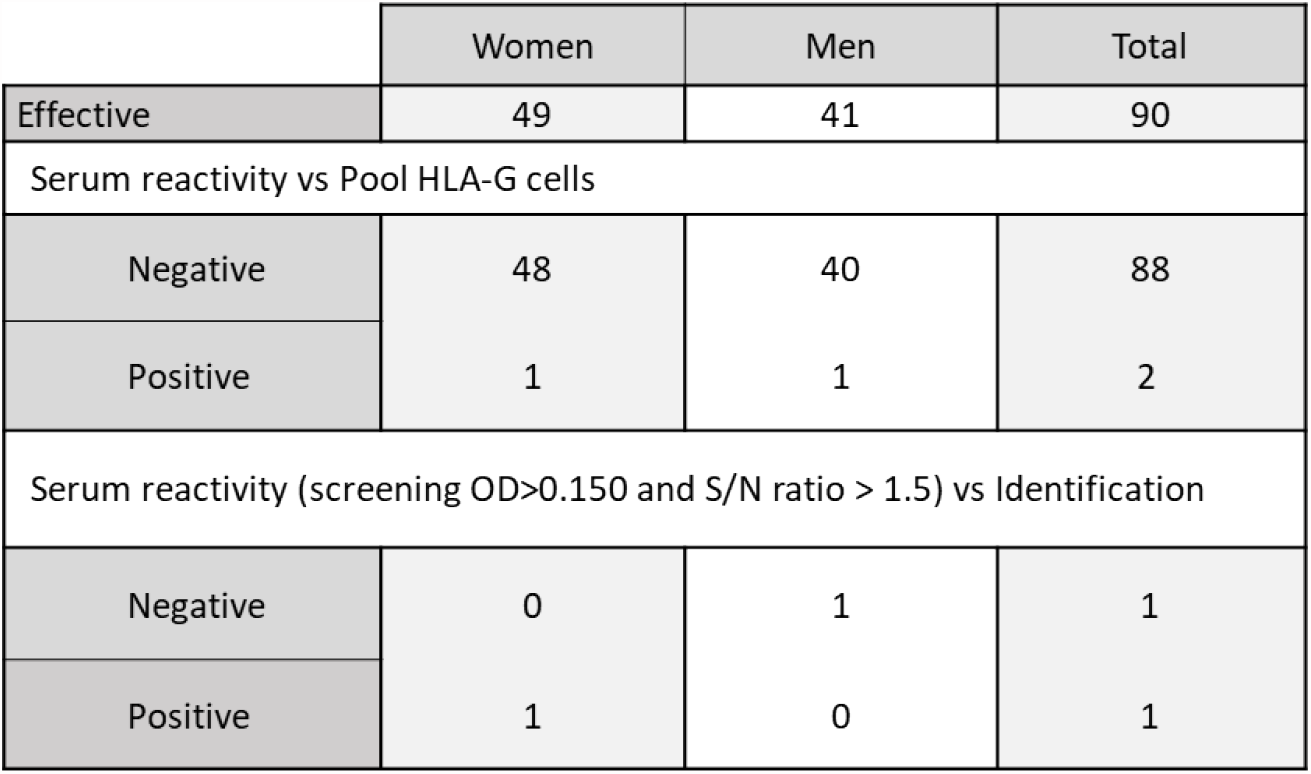
Results of adapted MAIPA on healthy subjects.

|  | Women | Men | Total |
| --- | --- | --- | --- |
| Effective | 49 | 41 | 90 |
| Serum reactivity vs Pool HLA-G cells |  |  |  |
| Negative | 48 | 40 | 88 |
| Positive | 1 | 1 | 2 |
| Serum reactivity (screening OD>0.150 and S/N ratio > 1.5) vs Identification |  |  |  |
| Negative | 0 | 1 | 1 |
| Positive | 1 | 0 | 1 |

### HLA-G antibody detection in Lung Transplantation Recipients

The pre-transplant sera (D0) and the post-transplant sera (from D15 to M12) from 35 LTRs were analyzed (Table 3). One LTR serum (ID=9) displayed positive results from the screening test at D15 and M1. However, these results could not be replicated for the two sera in the screening test, and no positive result could be obtained in the identification assay. HLA-G antibody detection was negative for all LTR sera.

**Table 3:**
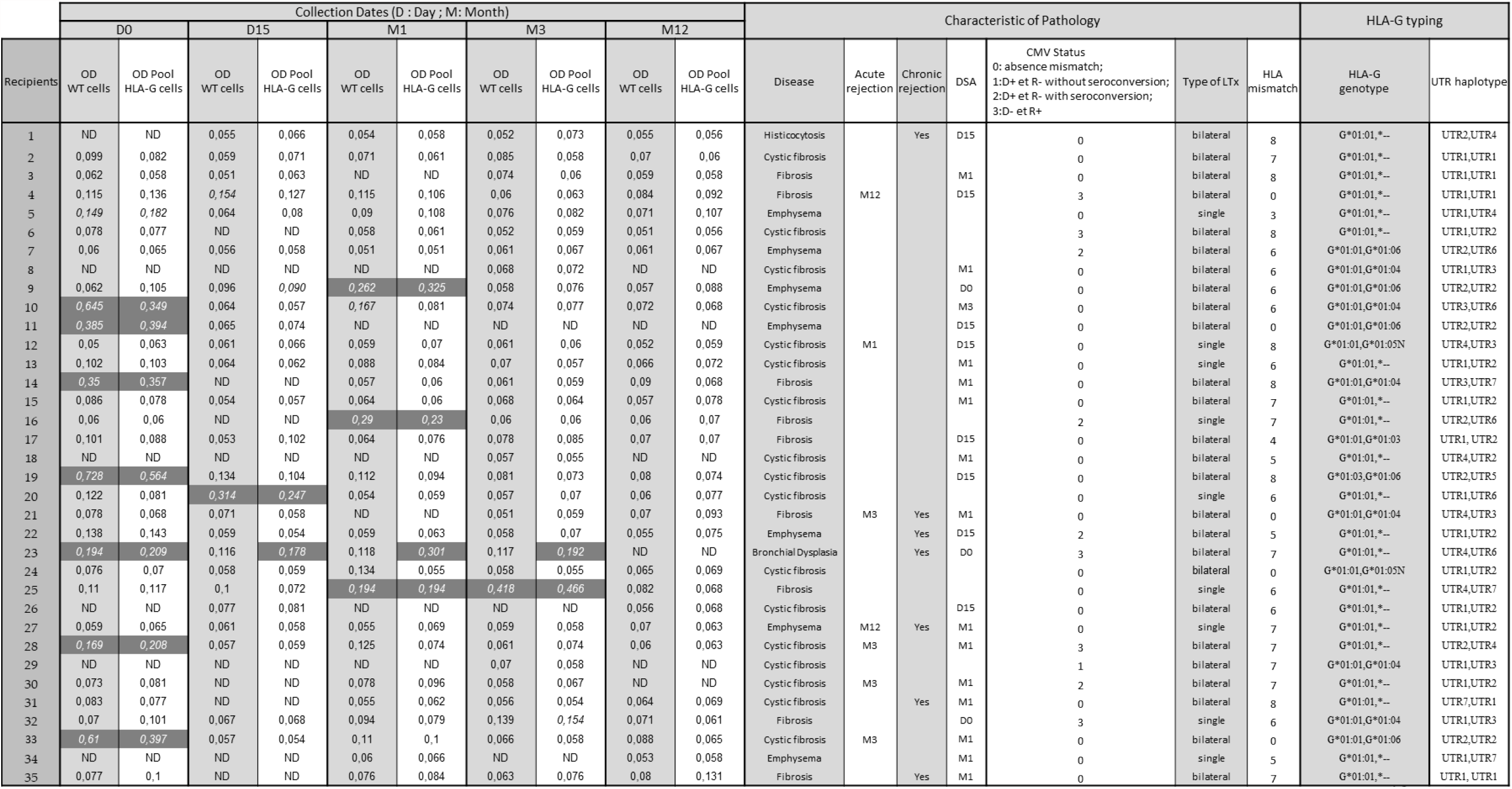
Results of adapted MAIPA in LTRs cohort at D0, D15, M1, M3 and M12 after LTx. Positive threshold: OD>0,150 (OD : optical density, WT : wild type, ND : not determined).

Sera from 11 LTRs had reactivity with an OD>150 for the HLA-G transduced lines pool, but with a ratio close to 1 between the HLA-G transduced and wild cell lines, signifying the detection of antibodies against an unidentified cell antigen expressed by the SPI-6 cell line. These cases were reproducible, either in the screening assay, identification assay, or both. This non-specific reactivity was detected in 7 sera before transplantation, in 1 serum at D15 after transplantation, in 5 sera at M1, and in 2 sera at M3. This non-specific reactivity was reproducible at different times in 3 LTRs (ID=23 at D0 and M1; ID=25 at M1 and M3; ID=9 at D15 and M1). This latter patient was transplanted for emphysema and produced DSA before LT, which persisted until M3 with a Mean Fluorescence Intensity (MFI) varying from 13,000 to 7,000. Patient ID=23 was transplanted for bronchial dysplasia. No humoral event was detected until M12. Patient ID=25 was transplanted for pulmonary fibrosis and produced DQ7 DSA with a stable MFI at 7,000 from D15 to M12. None of these 3 patients displayed acute rejection in the first year. Patient ID=23 reported chronic rejection. The blood groups of the three patients were different (data not shown). The data are summarized in Table 3 and in Supplementary table.

## DISCUSSION

Detection of HLA-G antibodies is poorly studied, although they may interfere with the immune regulation mechanisms that occur during and after organ transplant. In this study, we aimed to validate a protocol for detecting HLA-G antibodies in serum and to explore their impact on LTx outcomes. We used an adaptation assay from the MAIPA ApDIA® kit to analyze 234 sera from healthy donors and lung transplantation recipients. Surprisingly, the HLA control from MAIPA ApDIA ® kit was HLA-G reactive. This control is produced from plasma of HLA hyperimmunized donors, which origin is unknown. Interestingly, it is not reactive for HLA-E (data not shown), suggesting a HLA-G specificdetection. However, a cross-reactivity with epitopes from classical HLA should not be excluded.

Only 1 out of 234 sera was found to be reactive against all HLA-G molecules isoforms, suggesting the presence of HLA-G autoantibodies rather than alloantibodies. HLA-G antibodies were detected in one healthy donor, a 22-year-old woman (A+ blood group), without any risk factors for HLA alloimmunization, notably pregnancy. However, HLA-G antibodies were not detected in the 30 female healthy donors who had at least one pregnancy. Like classical HLA class I antibodies, HLA-G antibodies may gradually disappear only one month after delivery in first pregnancy and increase with the number of pregnancies. However, considering the crucial role of HLA-G in pregnancy success, HLA-G antibody may be more frequent in women who underwent pregnancy complications such as pre-eclampsia or unexplained fetal loss in the second and third trimester. Sera collected from pregnant women at the onset of complications and away from childbirth should be tested.

Thus, we could not confirm previous results, from Jucaud et al., that detected HLA-G antibodies in 6 out of 17 healthy subjects using Luminex method (19). Of note, our cohort size was greater. The adapted MAIPA assay used in the present study may be less sensitive than Luminex. However, classical HLA class I alloantibodies detection from platelet lysis by the MAIPA technique (ApDIA®) is as sensitive as the Luminex technique (data not shown). HLA-G antibody calibrated standard would allow sensitivity comparison.

Luminex is a very sensitive assay for HLA antibodies detection, as illustrated by HLA antibodies identification in non-alloimmunized healthy males (22). HLA cryptic epitopes may be exposed following antigens denaturation by β2m loss during the experiment process (23), as reported for HLA (24). The main assumption concerning the detection of these HLA antibodies without an immunizing phenomenon is a cross-reactivity with epitopes other molecules as bacterial or animal proteins or with non-classical HLA molecules (25).

Conversely, the adapted MAIPA assay detects the complete form of HLA-G associated with β2m. However, the lack of reactivity in our adapted MAIPA assay could be due to binding competition between the MEM-G/9 antibody and those present in the sera.

No HLA-G antibody was detected in sera collected at different times before and after transplantation from 35 LTRs, strongly suggesting that anti-HLA-G antibodies are not involved in the occurrence of lung transplant complications. Our hypothesis was to detect more HLA-G autoantibodies than HLA-G alloantibodies, although DSA detection was frequent in LTRs. Indeed, the occurrence of anti-HLA-G antibodies could be secondary to the increase in expression of HLA-G by the lung graft during an inflammatory syndrome. In our LTR cohort, only three patients had HLA-G phenotyping corresponding to low HLA-G expression. The UTR2, UTR5, and UTR7 haplotypes are associated with a decrease in cellular and soluble HLA-G expression (18). This weak expression can limit the production of anti-HLA-G antibodies directed against the graft. Similarly, only six donors had the haplotype of poor prognosis for the occurrence of chronic rejection. Furthermore, the HLA-G variability is low, and HLA-G antibodies have been detected in autoimmune diseases. However, in the latter cases, it cannot be completely excluded that this detection may rely on cross-reactivity with some epitopes expressed on HLA-E molecules or another non-classical and classical HLA. Finally, since data on the donor HLA-G status were missing, the confirmation of the alloimmunization process could also not have been considered.

Twelve LTRs (and one healthy donor) had non-specific reactivity against the non-transduced HLA-G cell lines following the MEM-G9 immunocapture. These results were accurate for three patients. One possibility is that the MEM-G9 monoclonal antibody can cross-reactivate with epitopes carried by classical HLA. However, this detection was not correlated with DSA detection, and the selected line did not express any HLA molecule. Whatever the reason for this increase in non-specific reactivity, the non-transfected line control is necessary.

Notwithstanding the limits of the adapted MAIPA assay, our results supported that the production of antibodies against the various non-denatured isoforms of HLA-G is rare in non-exposed individuals (healthy donors) and in alloimmunized patients (LTRs). Such results need to be confirmed by immuno-capture performed with other HLA-G monoclonal antibodies and with other multicentric cohorts.

## ABBREVIATIONS

(HLA): Human Leucocyte Antigen
(LTx): Lung Transplantation
(LTR): Lung Transplantation recipient
(Abs): Antibodies
(Ag): Antigen
(DSA): Donor Specific Antibodies
(BOS): Bronchiolitis Obliterans Syndrome
(D): Days
(M): Month
(MFI): Mean Fluorescence Intensity
(OD): optical density

## AUTHOR CONTRIBUTIONS

PP, LH supervised the study. JBB, AT performed the technical analyses. FJ, CB and AB participated in the technical expertise. FC, JC, MS and CP assisted in the interpretation of the results. M-RG and BC collected the clinical data from the patients. All authors contributed to the article and approved the submitted version.

## ACKNOWLEDGEMENTS

We thank Yasmina Denhadji and Ludovic Choucha, two technicians of Platelet Immunology department from EFS PACC, for their assistance and Dr Claude Bagnis for assisting in the construction of HLA-G cell lines.

## CONFLICTS OF INTEREST

The authors declare that the research was conducted in the absence of any commercial or financial relationships that could be construed as a potential conflict of interest.

## Figures

**Supplementary Figure 1:**
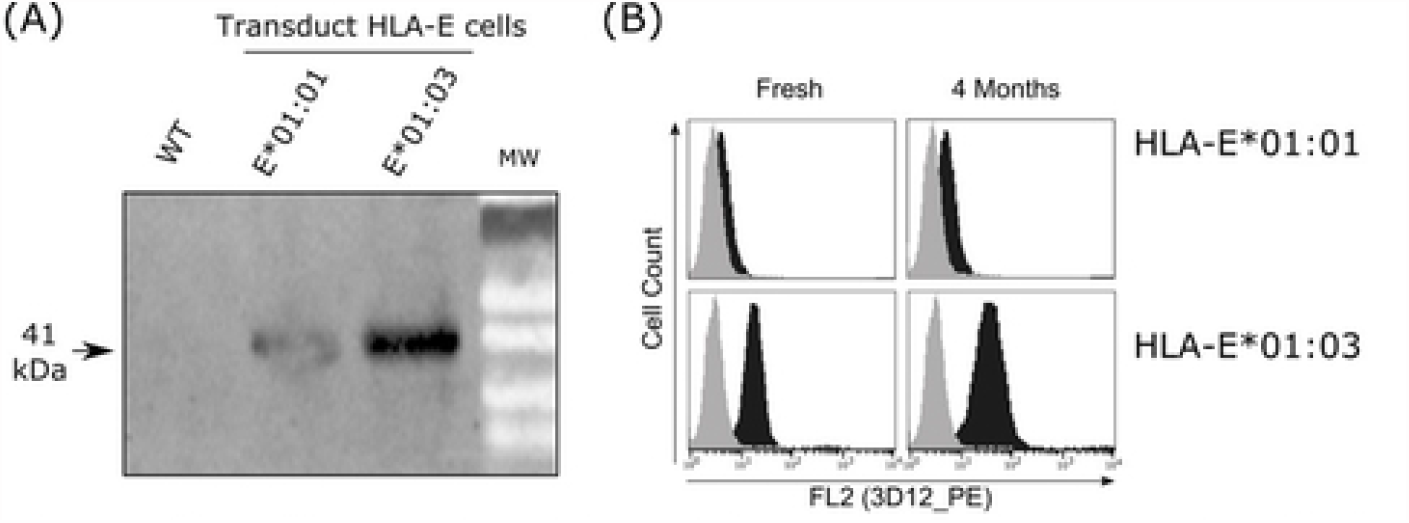
Expression of HLA-E isoform on SPI801 cell lines. (A) Western Blot analysis of HLA-E in the transduced cells in comparison with non-transduced cells (WT = wild type). (B) Stable expression levels of HLA-E isoform in transduced cells (black) versus non-transduced cells (gray) by flow cytometry.

**Supplementary figure 2:**
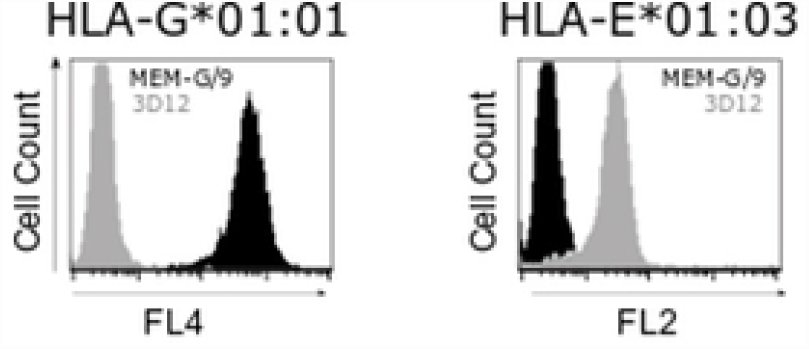
Specificity of detection of HLA-G and HLA-E by MEMG/9 and 3D12, respectively. Histograms show flow cytometry analysis of cells stained with MEM-G/9 (anti HLA-G antibody) in black or 3D12 (anti HLA-E antibody) in grey. Left histogram shows absence of detection of HLA-E on HLA-G*01:01 cells. Right histogram shows absence of detection of HLA-G on HLA-E*01:03 cells. Same analysis was performed on HLA-G*01:04, HLA-G*01:06 and HLA-E*01:01cells line (Data not shown).

**Supplementary Table:**
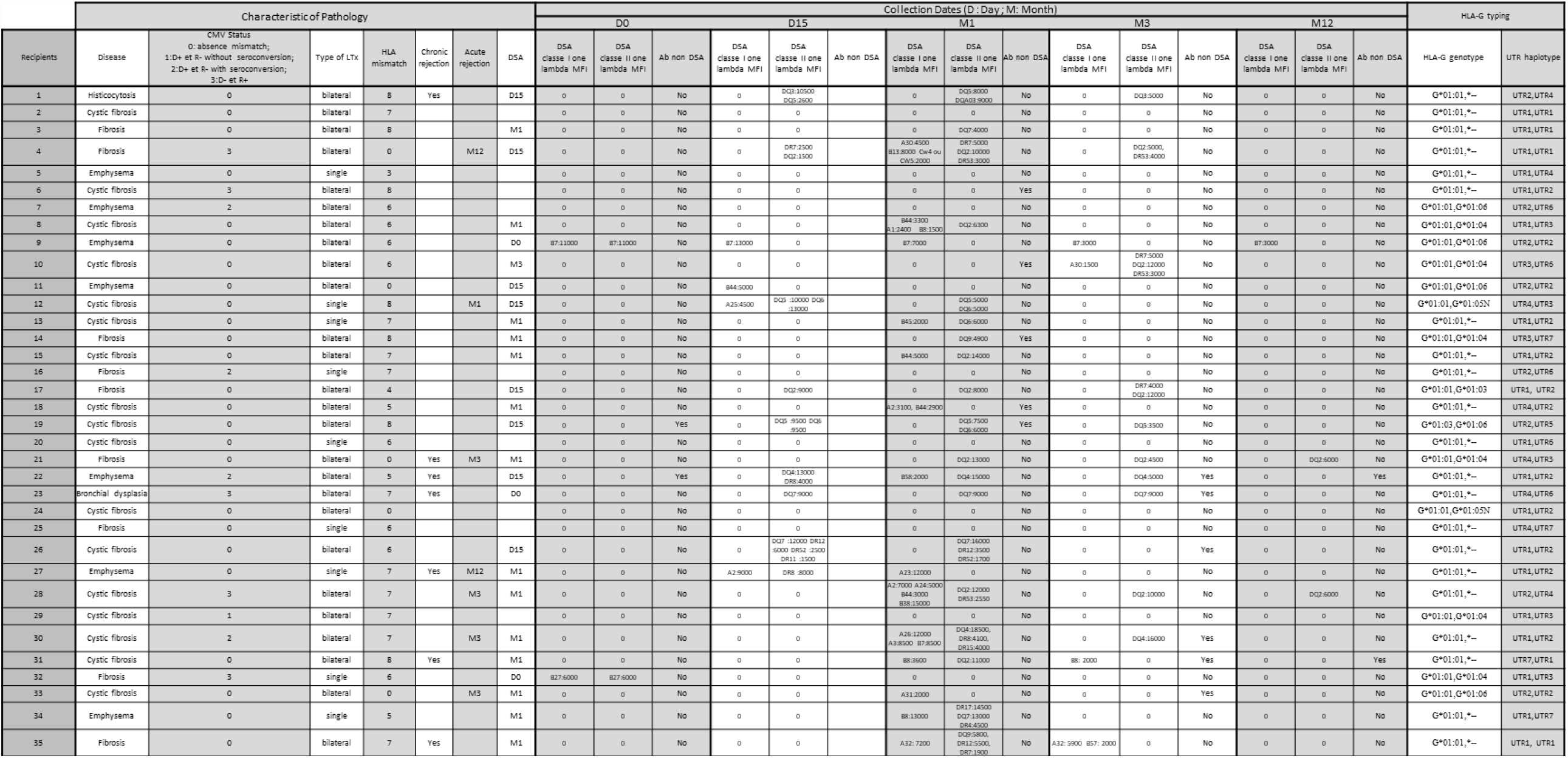
characteristics of the pathology, collection dates of the sera and results of the DSA research, and HLA-G typing results for each recipient

## Notes

### Competing Interest Statement

The authors have declared no competing interest.

